# Adaptation of Perceived Animacy from Biological Motion: Evidence for a “Life Motion Detector”

**DOI:** 10.1101/2025.09.03.673621

**Authors:** Mei Huang, Xinlin Yang, Geqing Yang, Li Shen, Zhaoqi Hu, Ying Wang, Yi Jiang

## Abstract

Humans can readily perceive animacy from biological motion (BM) - the distinctive movement patterns of living entities. However, how the human brain extracts animacy information from BM remains largely unclear. The current study investigated this issue using visual adaptation, a non-invasive tool for revealing neural mechanisms underlying the selective processing of specific properties. Results showed that prolonged exposure to intact human walkers, compared to non-BM adaptors, biased the perception of subsequent walking stimuli towards less animate. This adaptation aftereffect persisted following adaptation to local foot motions carrying diagnostic kinematic cues, but was absent after exposure to static body forms, indicating the involvement of neuronal populations encoding animacy from BM based on motion signals. Moreover, adapting to pigeon movements also biased animacy perception for human motions, revealing a shared mechanism for encoding animacy from cross-species BM signals. These results support the existence of an inherent “life detection” system in the human brain, attuned to local foot motions and kinematic features prevalent in vertebrate movements, which may lay a foundation for perceiving life from motion across terrestrial species.

**Public significance statements:** Discerning whether a moving entity is animate is crucial for survival and for generating appropriate responses. This study explored how the human brain perceives animacy based on a prevalent movement pattern among vertebrates, namely, walking. Leveraging a non-invasive visual adaptation paradigm, five experiments demonstrated that pre-exposure to intact human walking, feet-only walking, and even intact pigeon walking motion sequences - but not static human forms - elicited significant adaptation aftereffects in animacy perception of human walkers. These findings support the existence of neural substrates encoding perceived animacy based on biological motion signals, particularly those carried by local foot motions and cross-species kinematic features. Such mechanisms may be integral to an inherent ‘life detection’ system in the human brain.

## Introduction

Animacy perception, the process of perceiving entities as alive or possessing lifelike qualities, is essential for the survival and reproduction of animals (De Agrò et al.,2024; Lemaire & Vallortigara, 2023; Lorenzi & Vallortigara, 2021; Scholl & Tremoulet, 2000; Tremoulet & Feldman, 2000). Given the evolutionary significance of animate information, humans and some other animals are endowed with exquisite sensitivity to animate stimuli in the environment (such as the faces, motions, and sounds of living beings), and exhibit preferential attention to living entities relative to inanimate ones (Calvillo & Hawkins, 2016; Calvillo & Jackson, 2014; Di Giorgio et al., 2017, 2021; Gonan et al., 2024; G. Yang et al., 2024; New et al., 2007; Rosa-Salva et al., 2016). These findings raise the possibility that the perception of animacy is a fundamental part of human cognition that potentially engages specialized processing channels in the neural system.

The movements generated by humans and animals, termed biological motion (BM), are among the most ubiquitous and salient animate stimuli in the natural world. It delivers rich and vital cues for animacy perception. Besides carrying information about animals’ forms, BM conveys kinematic patterns that are unique to living organisms. Successful extraction of these motion cues facilitates rapid detection or discrimination of animate beings, even at long distances or in low visibility. However, despite its significance, how people perceive animacy from BM based on motion cues and whether this process engages specific neural substrates remains largely unclear.

Perceiving animacy from visual motion cues involves at least two distinct mechanisms, either mediated by high-level social cognitive processes or implemented through basic visual processing, as suggested by a recent review (Huang et al., 2023; Shen et al., 2025). The first mechanism enables observers to infer animacy (or agency) from movements conveying intention or interaction through social inferences (Csibra., 2008; Gao et al., 2010; Heider & Simmel, 1944; Meyerhoff et al., 2014). Observers irresistibly attribute social properties (e.g., intention and emotion) to simple geometric shapes interacting with each other or with the environment (e.g., chasing, avoiding, helping, hindering), generating an impression of agency that transcends mere animacy. The second mechanism, in contrast, focuses on the rapid, automatic perception of animate properties based on the visual processing of BM or simulated BM stimuli. Some studies simulate the external motion of an entity by imposing self-propelled motions on a simple object, evoking a sense of animacy through internal energy source metaphors (Scholl & Tremoulet, 2000; Stewart, 1983; Szego & Rutherford, 2008; Tremoulet & Feldman, 2000). However, since self-propelled motions can also be initiated by non-living things, such as a volcano or a spring, it can not fully explain our sensitivity to the distinctive movement patterns of living beings. To address this issue, other studies applied point-light BM stimuli that capture the intrinsic joint movements of humans or other vertebrate species (Chang & Troje, 2008; Johansson, 1973; Lorenzi et al., 2024; Lu et al., 2024; Thurman & Lu, 2013; Vallortigara et al., 2005). These stimuli reserve characteristic kinematic patterns arising from the coherent joint movements within animal bodies, the processing of which involves specialized mechanisms different from that for inanimate motions, as revealed by mounting behavioral, neuroimaging, developmental, and animal studies (Bardi et al., 2011; Carter & Pelphrey, 2006; Di Giorgio et al., 2017, 2021; Pavlova, 2012; Simion et al., 2008). Investigating animacy perception from these point-light BM stimuli is essential for understanding how humans identify living beings based on their unique movement patterns in real-world contexts.

Extensive evidence shows that people can extract a variety of properties (e.g., identity, walking direction, gender) from the point-light BM displays (Chang & Troje, 2008; Pollick et al., 2005; Y. Wang et al., 2022; X. Yang et al., 2014), and the extraction of some information can occur solely based on motion cues. Troje and Westhoff (2006) found that observers could discriminate walking direction even from spatially scrambled BM displays. Because spatial scrambling disrupts the global configuration but not the local kinematic cues, this result highlights the significance of local kinematic cues to walking direction perception. Moreover, scrambled and intact walking hens were equally effective in eliciting visual preferences in newborn chicks (Vallortigara et al., 2005). Based on these findings, researchers proposed that there is a visual filter in the human and animal brains attuned to the local motion of terrestrial vertebrates for detecting ‘life signals’ (Johnson, 2006; Troje & Westhoff, 2006; Vallortigara, 2012). Further research speculated that the ‘life signals’ might be primarily transmitted by the foot motions of walkers (Bardi et al., 2014; Chang & Troje, 2009a, 2009b), and such signals are detected by an innate ‘Step Detector’ rooted in the subcortical regions of the human brain (Hirai and Senju, 2020). However, note that the direction discrimination and preferential-looking tasks used in previous studies did not directly assess the perception of life, particularly to what extent observers perceived the motion as being animate. While one study required participants to rate the perceived animacy from spatially scrambled BM and found the rating scores correlated with the walking direction discrimination performance (Chang and Troje, 2008), it did not directly examine the role of local foot motions in animacy perception. Therefore, it remains to be elucidated whether the hypothesized life detection or step detection system contributes to the immediate perception of animacy from BM.

In addition to perceiving conspecific BM, humans can perceive the BM of other animals, such as horses, goats, baboons, elephants, and cats (Chang & Troje, 2008; Mather & West, 1993; Pinto & Shiffrar, 2009). Moreover, newly hatched chicks exhibited an innate preference for BM patterns regardless of whether the BM belonged to the same species or potential predators (Vallortigara et al., 2005). These results imply that the life-detection system may involve an evolutionarily ancient neural mechanism for detecting legged vertebrates in a broad range, attuned to cross-species motion signals despite their forms (Johnson, 2006; Troje & Westhoff, 2006; Vallortigara, 2012). It leads to a question of whether and to what extent cross-species movement cues could drive the life-detection system in the human brain and influence animacy perception.

The present study addressed the abovementioned questions using the visual adaptation paradigm. Repeated exposure to a stimulus with a given feature will cause the deviation of the observer’s perception of the following stimuli in the opposite direction. Such an adaptation aftereffect is considered to result from weakened neuronal responses selective to the tested feature of the adaptor, including high-level information such as the perceived animacy (Koldewyn et al., 2014). It thus provides a non-invasive approach to reveal the activity of neuronal populations underlying specific cognitive processes through behavioral performances (Clifford et al., 2007; Palumbo et al., 2017; Webster & Macleod, 2011). Combining the visual adaptation paradigm with psychophysical measurements, we conducted five experiments to unveil the mechanisms for animacy perception from BM cues. To test whether there exist neuronal substrates encoding the animate properties of BM based on motion cues, Experiments 1 and 2 investigated the animacy adaptation aftereffect induced by intact human BM adaptors while comparing it with the effect elicited by static BM configuration adaptors. Experiment 3 explored whether the adaptation effect in animacy perception can be driven by local foot motion signals, employing the movements of feet as adaptors. Experiments 4 & 5 further investigated whether and to what extent the adaptation aftereffect can transfer across species by adapting observers to intact human BM and pigeon BM, respectively, and comparing the animacy perception aftereffects.

## Method

### Participants

A total of 120 participants (ages ranged from 18 to 28 years, mean age ± SD = 23.28 ± 2.14 years, 74 female) took part in the study, with 24 participants in each of the five experiments (11 female in Experiment 1, 16 in Experiment 2, 16 in Experiments 3 and 4, and 14 in Experiment 5). All participants had normal or corrected to normal vision and were naive to the purpose of the experiments. All gave their written informed consent in accordance with procedure and protocols approved by the institutional review board before participating in the study. We determined the sample size using G*Power (Faul et al., 2007). A sample size of 19 participants would be sufficient (power = 0.90, α = 0.05) to detect a large animacy adaptation aftereffect (Cohen’s d > 0.8), based on a study using a similar paradigm to measure the adaptation aftereffect in face animacy perception (Koldewyn et al., 2014). The sample size was further increased to 24 per experiment to adequately detect the animacy adaptation aftereffect in the current study.

### Stimuli

The natural point-light BM sequence consists of fifteen luminous dots located on the head and key joints of a human walker, showing a realistic motion pattern of a person walking on a treadmill (Saunders et al., 2009; Troje, 2002). Each cycle lasts 1s and contains 30 frames. The non-BM sequences were created by distorting the motion trajectory of each light point of the BM stimulus and removing the acceleration cues along the modified trajectory (Troje & Chang, 2013). The modified trajectory was either the circumcircle of the original natural trajectory (in Experiments 1-3) or along a horizontal line segment passing through the center of gravity of the original trajectory (in Experiments 4-5). We adopted these two different methods to destroy the special kinematic feature of BM in order to control for the potential effect of specific stimuli on the results.

The adapting stimuli were varied across experiments to examine the roles of different BM cues in the adaptation aftereffect (**Fig.1b**, adaptors). In Experiments 1 and 4, the intact natural BM sequence and the non-BM sequence (approximately 2.0°× 7.4° in visual angle) were employed as the adaptors to investigate the overall adaptation aftereffect induced by the BM cues. In Experiment 2, to explore the role of the global form, the adaptors were static frames extracted from the BM and the non-BM sequences used in Experiment 1. For each sequence, we extracted two frames that represented static forms with the maximal horizontal and vertical extensions, respectively, within a motion cycle. Experiment 3 was aimed to test whether foot movements can drive the perception of animacy, thus a feet motion sequence with two dots at the ankles and its inverted non-BM version (subtended approximately 2.7°×0.29° in visual angle) served as adaptors. In Experiment 5, the adaptors were point-light pigeon BM, which had two light dots attached to each foot and two light dots on the head and the beak, and the inverted non-BM version of the pigeon BM.

**Figure 1.**
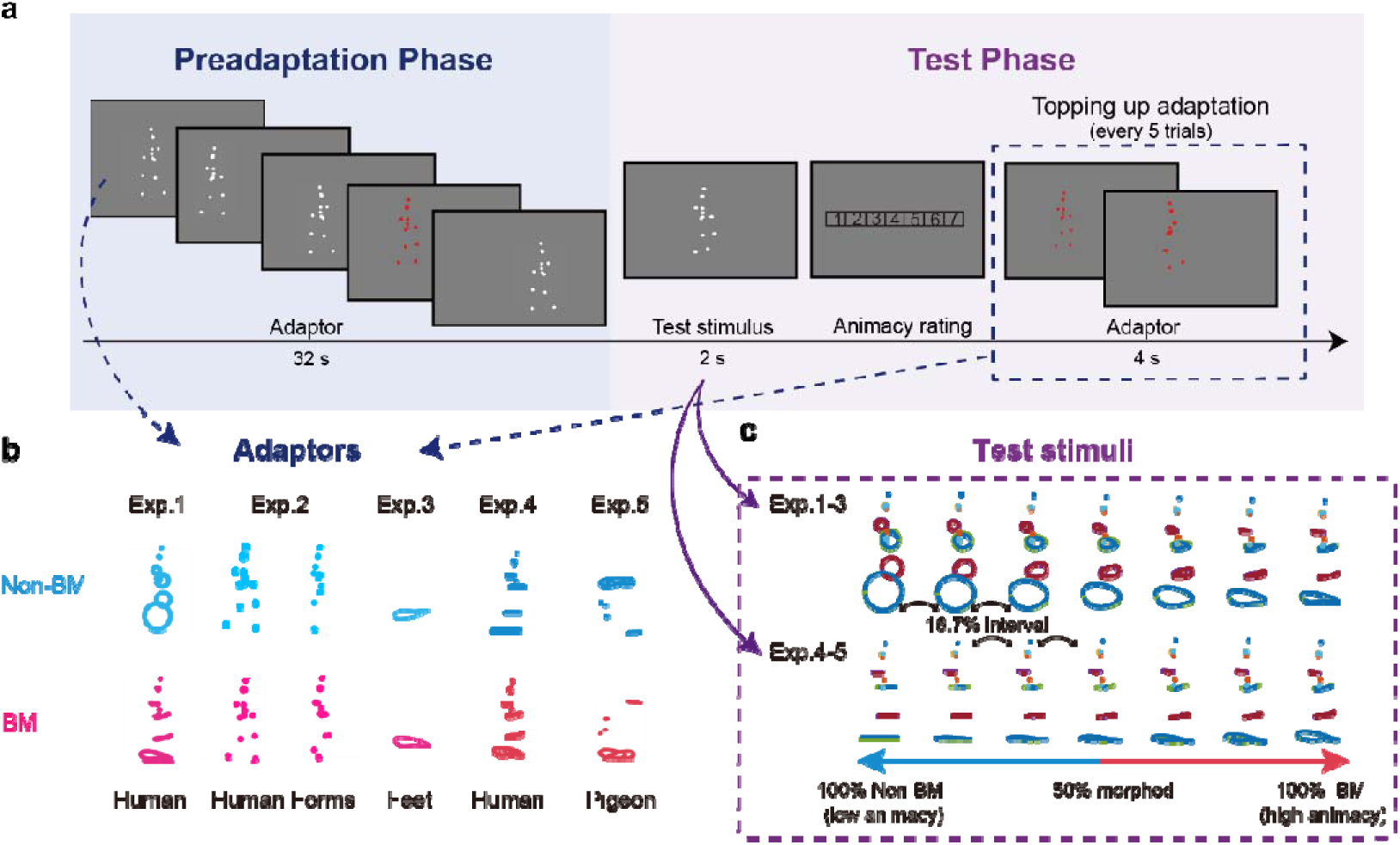
Illustrations of the experimental procedure for Experiment 1 (a) and the adaptors (b) & test stimuli (c) used in Experiments 1–5. **a.** The experiment started with a preadaptation phase, during which a biological motion (BM) or a non-biological motion (non-BM) adaptor was presented for 32 s. Participants were required to watch the movement of the stimulus and detect color changes that occurred every 3-5 s. During the test phase, participants were required to rate the animacy level of a motion-morphing stimulus on a 1-7 scale in each trial. There was a 4-s topping-up adaptation session before every five trials to maintain the adaptation effect. **b.** The motion trajectories of each dot of the adaptors used in the non-BM and BM conditions in each experiment. **c.** The motion trajectories of each dot of the test stimuli at seven morph levels in each experiment. The colors of the dots are for illustration only. The stimuli were rendered in white in the experiments.

In each experiment, a set of test stimuli was generated by morphing the motion trajectories between the natural BM (the most animate) and the non-BM stimuli (the least animate) at seven levels (**Fig.1c**). Each test stimulus was generated with a certain weight for both the BM and non-BM sequences, where the weights of BM stimuli were 0,0.167,0.333,0.5,0.667,0.833,1 in order, and the weights of non-BM stimuli were 1, 0.833, 0.557, 0.5, 0.333.0.167, 0. The non-BM sequences were different between Experiments 1-3 and 4-5 as mentioned above to assess the generalizability of the results.

The point-light motion stimuli were displayed using MATLAB (Mathworks, Inc., Natick, MA) together with the Psychtoolbox extensions (Brainard, 1997; Pelli, 1997). The stimuli were rendered in white against a uniform grey background and displayed on a 24-inch LCD monitor (1920×1080, 60Hz) with a viewing distance of 60 cm. The initial frame of the point-light display was randomized for each trial.

### Procedure

**Experiment 1.** In a practice session, participants were required to rate the perceived animacy for the 7 morphed point-light motion displays presented in a randomized order. Each sequence was presented for 2000 ms, and participants rated each stimulus 10 times, at a 1-7 scale, where 1 means that the stimulus has almost no animacy, and 7 means that the stimulus has the highest level of animacy. They were told that the criteria for rating animacy is the extent to which the display is a true and natural reflection of the movement of living things on Earth. The purpose of practice was to ensure that the participants got familiar with the task and the differences among the seven morphed motion stimuli.

The formal experiment included two conditions: adaptation to a normal intact BM and adaptation to an acceleration-deprived non-BM. The order of these conditions was counterbalanced across participants. Each condition comprised both a preadaptation phase and a test phase (**Fig. 1a**). During the 32s preadaptation phase, a normal BM or a non-BM sequence was presented. To eliminate the effect related to the adaptation of low-level visual properties, the stimuli’s facing direction was changed every 2 seconds, and the center location of the stimuli was jittered randomly within a square area of 1.3°×1.3°. The color of the point-light stimuli changed to red every 3-5 seconds. Participants were required to view the movement of the stimulus and press the space key immediately when they detected the color change. The preadaptation phase was immediately followed by a test phase in which participants judged the animacy of a series of motion stimuli. The stimuli consisted of seven morph levels, each of which appeared 20 times, half facing left and half facing right, resulting in a total of 140 trials presented in random order. Every 5 rating trials were preceded by a 4s ‘topping-up’ adapting stimulus (same as the adaptor in the preadaptation phase but always in red) to maintain the adaptation effect.

**Experiment 2.** The preadaptation phase was identical to Experiment 1, except that the adaptors were static frames extracted from the BM and non-BM sequences. During the 32-second preadaptation, two static frames (maximal horizontal/vertical extensions) alternated every 2 seconds. The topping-up adaptation stimuli also contained two different static frames alternated in a random order. The test phase remained identical to Experiment 1, with participants rating the same seven morphed motion stimuli.

**Experiment 3.** The design matched Experiment 1, but adaptors were simplified to two-dot motion sequences depicting the upright foot movements (BM) and the inverted counterparts (non-BM). During preadaptation, participants viewed foot motion sequences with randomized direction/location jitter. The test phase was the same as previous experiments.

**Experiment 4.** Experiment 4 replicated Experiment 1 in structure but applied a different method to generate non-BM stimuli (see Stimuli for details). The adaptors and test stimuli were updated accordingly, while the procedure remained unchanged.

**Experiment 5.** Experiment 5 differed from Experiment 4 in that it employed upright walking pigeons as the BM adaptors and the inverted pigeons as the non-BM adaptors. Test stimuli remained human BM and non-BM morphed sequences. Participants underwent the same preadaptation and test phases. They were unaware of the pigeon form, as confirmed by post-experiment inquiries.

### Data Analysis

First, the 1-7 rating scores for perceived animacy were transformed to a 0-1 scale with min-max normalization for each participant. The normalized scores from each participant for each adaptation condition were fitted with a sigmoid Boltzmann function f(x)=1/(1+exp((x–x0)/ω)), where x represents the morph level of the test stimuli from non-BM to BM (0,0.167,0.333,0.5,0.667,0.833,1). We estimated the point of subjective equality (PSE), or x0 at which the observer perceived a motion sequence as equally animate and inanimate, and the difference limen (DL), an index of animacy discrimination sensitivity corresponding to half the interquartile range of the fitted function. The PSE and DL values were then subjected to further statistical analyses. A PSE shift toward a larger value after adapting to BM relative to non-BM cues, with no change in DL, indicates the presence of an adaptation aftereffect, that is, adaptation to BM cues biases the subsequent animacy perception toward less animate.

### Transparency and openness

The present study has reported how the sample size was determined, as well as all manipulations and measures used in the study. This study followed JARS (Kazak, 2018) and was not preregistered. All data are available at https://osf.io/qgz53/?view_only=371981924e914cf2bdf012353d1e83d6. Data were analyzed using MATLAB (Mathworks, Inc., Natick, MA) and JASP, version 0.16.40.

## Results

The performance of the color detection task was higher than 95% in all experiments (95%, 97%, 95%, 98%, 97% in Experiments 1-5), indicating that participants had directed their attention to the adaptors throughout the preadaptation phase. The data from the test phase was then analyzed to assess the adaptation aftereffect.

Experiment 1 investigated whether adapting to natural human BM cues could lead to an adaption aftereffect in animacy perception. As expected, a paired-sample t-test revealed a significantly larger PSE in the adapting BM condition compared to the adapting non-BM condition (**Fig.2a**, t(23) = 4.63, p < 0.001, Cohen’s d = 0.945, 95% CI for the mean difference = [0.035, 0.093], BF_10_ = 235), suggesting that the test motion was perceived as less animate in the former condition than in the latter condition. Furthermore, there were no significant differences in participants’ animacy discrimination sensitivity (i.e., DL) between the adapting BM and adapting non-BM conditions (t(23) = -1.058, p = 0.301, Cohen’s d = -0.216, 95% CI for the mean difference = [-0.064, 0.021], BF_10_ = 0.354). These results clearly demonstrated that preexposure to intact human BM cues induced significant adaptation aftereffects on animacy perception.

**Figure 2.**
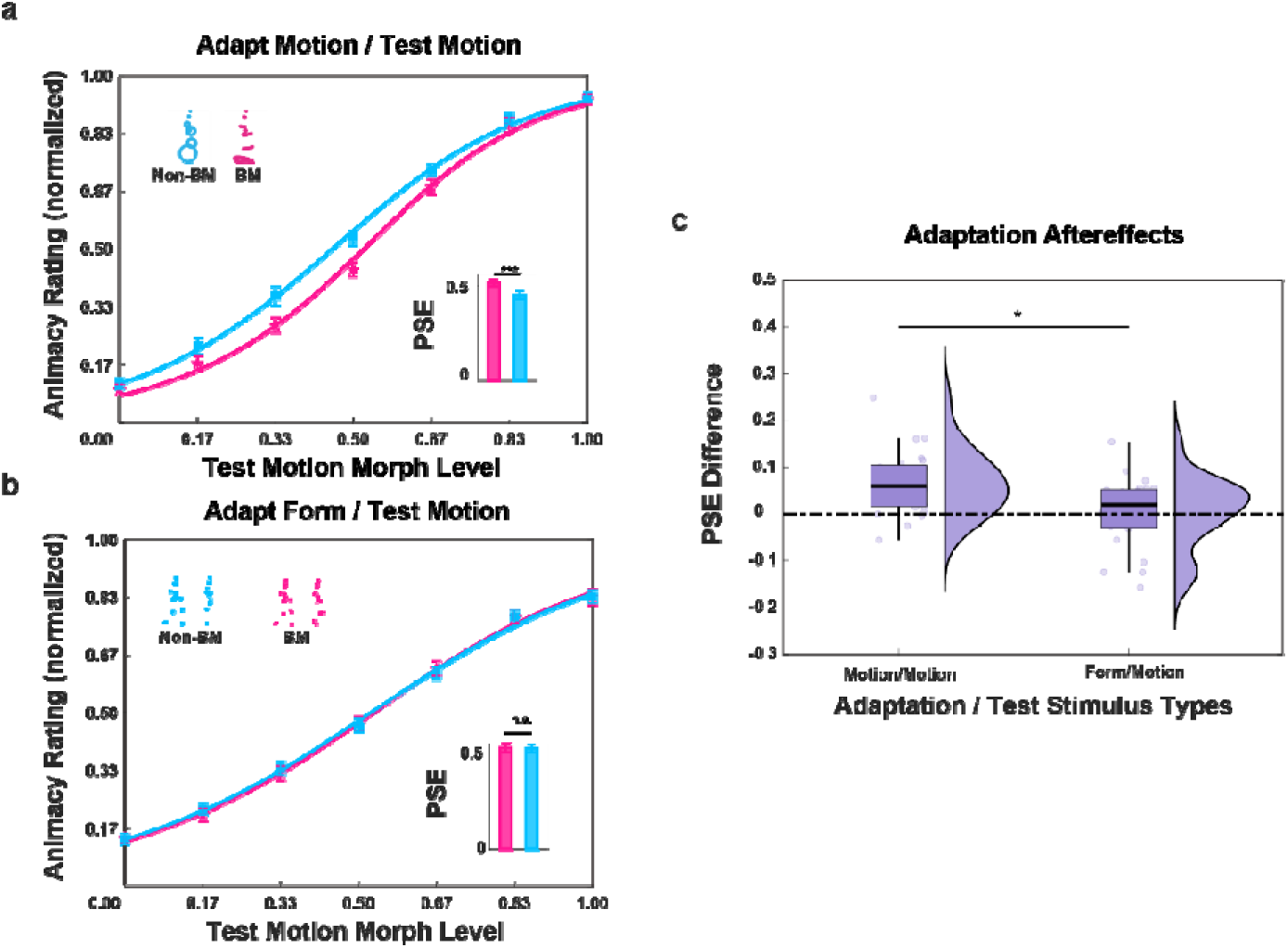
Results for Experiments 1-2 (a, b) and the comparison of the adaptation aftereffects across experiments (c). Pink and blue lines represent psychometric curves derived from data obtained from the adaption to BM and non-BM (**a**) or BM and non-BM forms (**b**) conditions. The inset bar charts show the PSEs for the two adaptation conditions. The error bars represent standard errors of means. *p < .05; ***p < .001. n.s., not significant.

### Experiment 2

Since natural human BM sequences convey both form and motion cues, and there are apparent differences in the global forms of the BM and non-BM stimuli besides their motion patterns in Experiment 1 (**Fig. 1**), it remains unclear whether the observed effect was derived mainly from the adaptation to motion signals. To solve this issue, Experiment 2 examined the aftereffect on animacy perception when subjects adapted to static forms of the human BM and non-BM stimuli. A paired-sample t-test revealed no significant difference in PSE between the two adaptation conditions (**Fig.2b,** t(23) = 0.258, p = 0.799, Cohen’s d = 0.053, 95% CI for the mean difference = [-0.028, 0.035], BF_10_ = 0.221), and the DLs were not significantly different between the two conditions (t(23) = -0.673, p = 0.508, Cohen’s d = -0.137, 95% CI for the mean difference = [-0.059, 0.030], BF_10_ = 0.264), indicating that preexposure to static forms of the human BM and non-BM stimuli could not elicit an adaptation aftereffect as in Experiment 1. Moreover, a comparison of the observed adaptation aftereffects (PSE differences) between Experiment 2 and Experiment 1 using an independent samples t-test revealed that the aftereffect in Experiment 1 was significantly higher than that in Experiment 2 (**Fig.2c,** t(46) = 2.929, p = 0.005, Cohen’s d = 0.846, 95% CI for the mean difference = [0.019, 0.102], BF_10_ = 8.056). These results rule out the confounding effect of stimulus forms in Experiment 1, suggesting that the observed aftereffect is more likely to result from the adaptation to visual motion cues.

### Experiment 3

In Experiment 3, we further investigated whether local motion signals in foot movements play a critical role in animacy perception from BM. Similar to Experiment 1, a significant increase of PSE was observed in the adapting to BM (upright feet motion) condition as compared to the adapting to non-BM (inverted feet motion) condition (**Fig.3a,** t(23)=2.235, p=0.035, Cohen’s d=0.456, 95% CI for the mean difference = [0.002, 0.062], BF_10_=1.711), suggesting that after adapting to the local foot motion cues, participants tended to perceive the test stimuli as less animate. In addition, the participants’ DLs did not show significant differences between the two conditions (t(23)=0.402, p=0.692, Cohen’s d=0.082, 95% CI for the mean difference = [-0.034, 0.050], BF_10_=0.231). Intriguingly, while the adaptation aftereffects (i.e., PSE differences) tended to decrease in Experiment 3 when compared to Experiment 1, the differences did not reach significance (**Fig.3b,** t(46) = -1.584, p = 0.120, Cohen’s d = 0.457, 95% CI for the mean difference = [-0.009, 0.0772], BF_10_ = 0.791), suggesting that the local motion of the feet is a primary cue that drives the animacy perception mechanism in the human brain.

**Figure 3.**
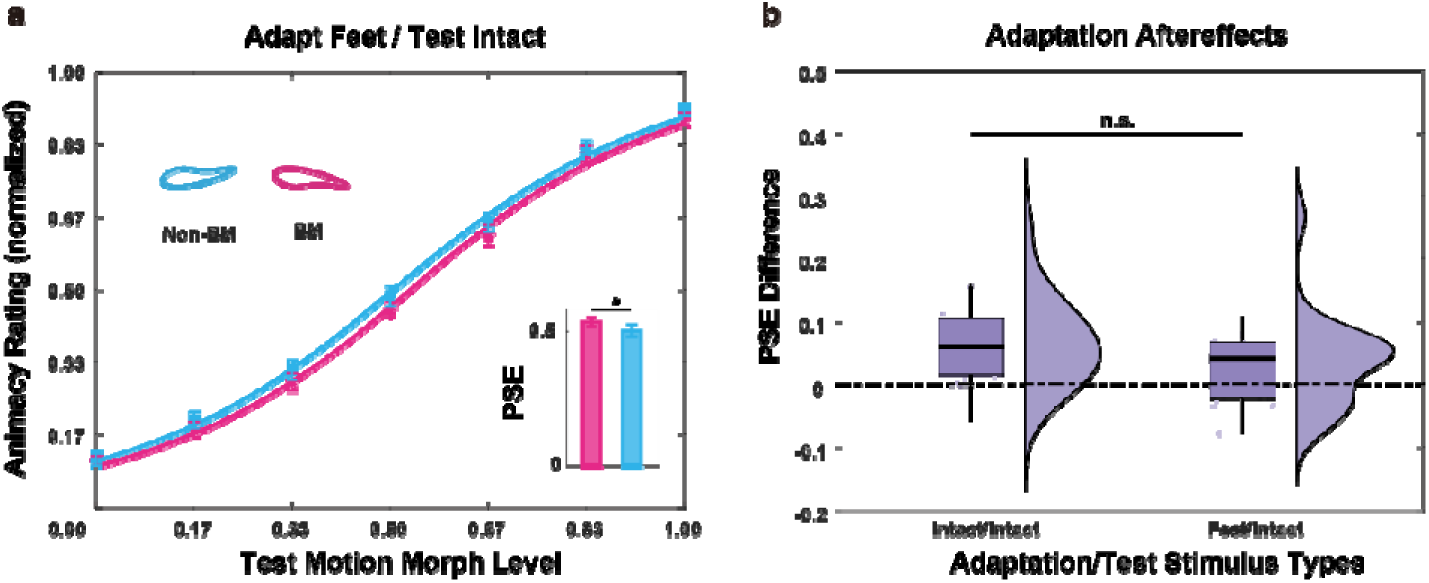
Results for Experiment 3 (a) and its comparison with the adaptation aftereffect observed in Experiment 1 (b). Pink and blue lines represent psychometric curves derived from data obtained from the adaptation to BM and non-BM (feet only) conditions. The data fitted by the psychometric function were plotted using solid lines. The inset bar charts show the PSEs for the two adaptation conditions. The error bars represent standard errors of means. *p < .05; ***p < .001. n.s., not significant.

### Experiment 4

To verify that the adaptation effect of animacy perception observed in the aforementioned experiments was not restricted to the given stimuli, we employed a new approach to generate another reconstructed non-BM sequence, adopting it as the non-BM adaptor and the corresponding morphed stimuli as the test stimuli in Experiment 4 (see Method for details). A significantly larger PSE was found in the adapting human BM than in the adapting reconstructed non-BM condition (**Fig.4a,** t(23)=3.816, p<0.001, 95% CI for the mean difference =[0.029,0.098], Cohen’s d=0.779, BF_10_=39.0). Meanwhile, there were no significant differences in DL between the two conditions (t(23)=-1.064, p= 0.298, 95% CI for the mean difference =[-0.041,0.013], Cohen’s d=-0.217, BF_10_=0.356). These results were in line with the findings from Experiment 1, eliminating the possibility that the given stimuli led to the results obtained from the previous experiments.

**Figure 4.**
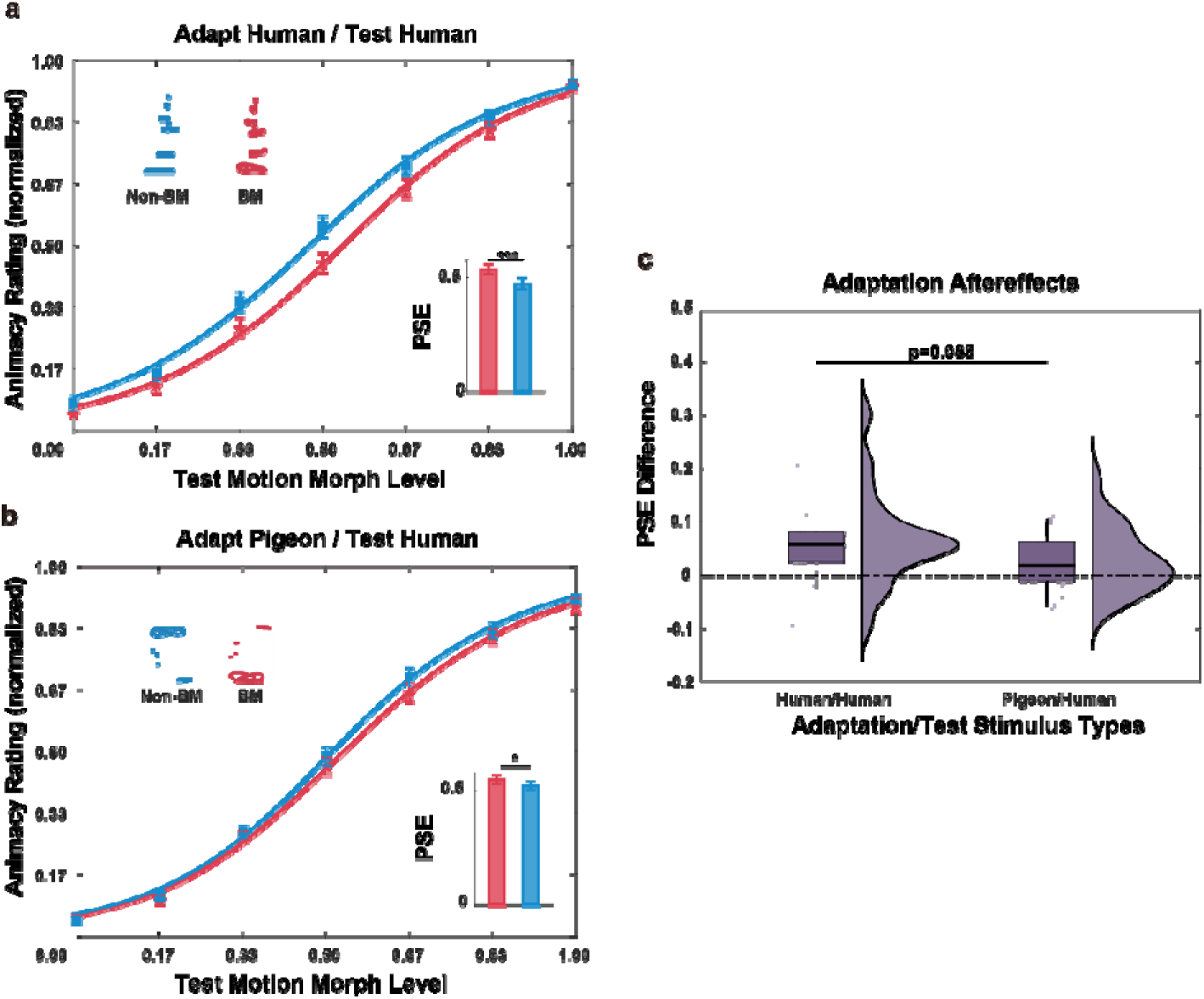
Results for Experiments 4 (a) and 5 (b), and the comparison of PSE difference between Experiments 4 and 5 (c). Red and blue lines represent psychometric curves derived from data obtained from the adaptation to BM and non-BM (**a**: human; **b**: pigeon) conditions. The inset bar charts show the PSEs for the two adaptation conditions. The error bars represent standard errors of means. *p < .05; ***p < .001.

### Experiment 5

To further investigate whether animacy perception from BM can be modulated by cross-species signals, in Experiment 5, the adapting stimuli were changed to a point-light walking pigeon and its non-BM control (i.e., inverted pigeon). Results showed that adapting to the pigeon BM could trigger an animacy adaptation effect compared to its non-BM control with a significant difference in PSE (**Fig.4b,** t(23) = 2.355, p = 0.027, 95%CI for the mean difference = [0.003,0.052], Cohen’s d=0.481, BF_10_=2.105) and no evident difference in DL (t(23) = 0.947, p = 0.354, 95%CI for the mean difference = [-0.021,0.055], Cohen’s d = 0.193, BF_10_ = 0.321). These results indicate that the mechanism underlying animacy perception from BM cues is sensitive to cross-species BM signals. Notably, inquiry after the experiment showed that most subjects were unaware of the fact that the adapting stimuli contained the form of pigeons, suggesting that the observed effect likely resulted from the adaptation to motion cues regardless of the stimulus form. Moreover, the adaptation aftereffect was smaller in Experiment 5 (adapting pigeon motion) than in Experiment 4 (adapting intact human motion) with a marginal significance (**Fig.4c,** t(46) = -1.761, p=0.085, 95% CI for the mean difference = [-0.005, 0.077], Cohen’s d=0.508, BF_10_ = 0.999), suggesting that although the animacy perception mechanisms are attuned to cross-species motion signals, such an effect is not as strong as that elicited by the same-species signals.

## Discussion

The present study investigated how the human cognitive system achieves animacy perception based on the motion cues in BM through the visual adaptation paradigm. Regarded as the psychologist’s microelectrode, visual adaptation provides a potent tool to probe the temporary reduction in activity of specific neuronal populations responsive to given attributes, offering insights into the functional properties and the neural mechanisms of visual perception (Webster & Macleod, 2011). Leveraging this tool, we found that pre-exposure to intact natural BM sequences (Experiments 1 & 4) induced an adaptation aftereffect on animacy perception, as revealed by a significant shift of perceived animacy of morphed human-like motions toward the opposite direction of the adaptor. These findings lend support to the existence of neuronal substrates in the human brain underlying the perception of animacy from walking—a pervasive and fundamental mode of human locomotion. Notably, such an aftereffect did not occur after adapting to static forms of the point-light walkers (Experiment 2), suggesting that it may be triggered by the kinematic cues in locomotion, which are generated through combined actions of bones and muscles under the constraints of gravity (Y. Wang et al., 2022), in a way highly distinct from the movement of non-biological entities.

Further results showed that prolonged viewing of local foot motions alone could induce a significant adaptation aftereffect (Experiment 3), underscoring the critical role of local motion cues, particularly those emanating from the feet, in animacy perception. Previous research suggests that scrambled BM conveying only local kinematic cues is still effective in eliciting visual preferences toward animate motions (Vallortigara et al., 2005) and leads to a perception of animacy (Chang and Troje, 2008). Among the local motions of different body joints, foot motions contain diagnostic cues that are critical to walking direction discrimination and presumably contribute to life detection (Troje & Westhoff, 2006). Hirai and Senju (2020) proposed a two-process theory of BM processing, wherein the first stage engages a ‘Step Detector’ system optimized for rapid processing of local foot motions and feet-below-the-body information. This system may facilitate the orienting to BM through an innate mechanism. While these theoretical accounts hint at the possibility that the foot motions of vertebrates are essential for perceiving animate information from motion cues, there is still a dearth of empirical evidence. Beyond the existing findings supporting the significance of foot motions in walking direction discrimination and attentional processing of BM (Bardi et al., 2014; Chang & Troje, 2009b, 2009a; Hirai et al., 2011; Shen et al., 2023; L. Wang et al., 2010; Y. Wang et al., 2018), the present study provides more direct evidence for the existence of a life detection mechanism in the human neural system attuned to foot motions, highlighting its pivotal role in animacy perception.

Moreover, adapting to pigeon movements could also bias the animacy perception of human BM, although to a slightly weakened degree than adapting to human movements (Experiments 4 & 5). These findings reveal that the neural encoding of perceived animacy from BM can transfer to some extent across species, which is in line with the assumptions about the pervasiveness of the ‘Life Detection’ or ‘Step Detector’ system across vertebrates (Hirai & Senju, 2020; Simion et al., 2008; Troje & Chang, 2023; Rosa Salva et al., 2015; Vallortigara et al., 2005; Vallortigara & Regolin, 2006). More crucially, these findings imply that the function of these systems may involve the discrimination of animate motion signals from inanimate motion.

Notably, this result contrasts with the previous finding that animacy perception from faces does not transfer between different species, specifically humans and dogs (Koldewyn et al., 2014), implicating potential disparities in animacy perception from BM and faces. Similar to BM, face-like stimuli elicit early visual preferences in several species (Kobylkov et al., 2024; Kobylkov & Vallortigara, 2024; Sugita, 2008). However, a monkey study showed that although such an innate visual sensitivity could generalize to human and monkey faces in infant monkeys reared with no exposure to facial stimuli, it developed into a species-specific effect after facial exposure during the sensitive period (Sugita, 2008). This narrowing process reflects the development of a general face prototype into a concrete representation, potentially engaging an attentional mechanism for the processing of socially meaningful, familiar faces. For BM, however, the innate cross-species preference towards its kinematics may not develop into a narrowed sensitivity even after visual exposure to conspecific motions, as suggested by the current findings, probably due to the crucial role of BM signals in life detection. The discrepancies in the cross-species properties of BM and face perception may stem from the differences in these stimuli regarding their evolutionary and social significance, as well as the distinction between the developmental courses of form- and motion-based animacy processing. Whether face- and BM-based animacy perception engages overlapping or distinct neural mechanisms remains an open question for future research.

Previous research implicates posterior superior temporal sulcus (pSTS) and intraparietal sulcus (IPS) in processing animate motions (Beauchamp et al., 2003; Duarte et al., 2022; Landsiedel et al., 2022; Pelphrey et al., 2003, 2004; Saygin et al., 2004; Schultz & Bülthoff, 2019; Sokolov et al., 2018; Walbrin et al., 2018). In parallel, neuroimaging studies on humans and monkeys suggest that some subcortical regions, including the superior colliculus (SC) and the ventral lateral nucleus (VLN), may be involved in the ‘Life Detector’ and ‘Step Detector’ system responding to local BM cues (Chang et al., 2018; Hirai & Senju, 2020; Lu et al., 2024;,Troje & Chang, 2013). Complementing this evidence, studies in newly hatched chicks have shown that subcortical regions including the preoptic area of the hypothalamus, the nucleus taeniae of the amygdala, and the right septum exhibit selective activation in response to BM (Lorenzi et al., 2024; Mayer et al., 2017). Together, these findings suggest that subcortical nuclei may constitute an evolutionarily conserved neural substrate for detecting life motion signals across species. However, the precise role of these subcortical regions and their connections with the cortical regions in animacy perception remains poorly understood. Further research could explore whether some shared locomotion characteristics among vertebrates, especially those embedded in local foot movements, instigate animacy perception in humans via an integrated subcortical-cortical network (Huang et al., 2023).

## Constraints on Generality

The adaptation aftereffect observed can be elicited by the walking movements of both humans and pigeons. However, questions remain regarding the generalizability of these findings across different types of motion and species. Compared with terrestrial animals, animals in air or water exhibit different movement patterns (e.g., swimming and flying) under the constraints of different environmental forces (Han et al., 2023; Larsch & Baier, 2018; Ma et al., 2022). Therefore, involving more species and motion types in further investigation may help precisely delineate to what extent the animacy perception mechanism is sensitive to heterospecific BM signals. In addition, all participants in the current study were young college students around 20-30 years old, which may pose limitations when generalizing these results to a more diverse group. Including children or infants as the subject group could provide further evidence to support the claim that the “life-detector” system in the human brain is innate or even genetically driven.

## Conclusion

The present study found significant adaptation aftereffects in animacy perception induced by intact human BM, human foot movements, and pigeon BM, but not when static human forms served as adaptors. These findings together provide converging evidence that perceiving animacy from BM involves a neural mechanism driven by local foot motions and responsive to cross-species kinematic cues, which may be integral to a ‘life detection’ system in the human brain.

## Acknowledgment

We thank Nikolaus Troje for kindly providing us with the point-light biological motion stimuli. This research was supported by grants from the Ministry of Science and Technology of China (2021ZD0203800, 2021ZD0204200), the National Natural Science Foundation of China (32171059, 32430043), the Interdisciplinary Innovation Team (JCTD-2021-06), the Youth Innovation Promotion Association of the Chinese Academy of Sciences, and the Fundamental Research Funds for the Central Universities.

## Author contribution

Mei Huang: Methodology, Software, Formal analysis, Investigation, Data curation, Visualization, Writing-original draft.

Xinlin Yang: Investigation, Writing-review and editing. Geqing Yang: Writing-review and editing.

Li Shen: Writing-review and editing.

Zhaoqi Hu: Methodology, Writing-review and editing.

Ying Wang: Conceptualization, Methodology, Supervision, Writing-review and editing, Funding acquisition.

Yi Jiang: Writing-review and editing, Funding acquisition.

## Data Availability

Data and stimuli are publicly available on the Institutional Knowledge Repository, Institute of Psychology, Chinese Academy of Sciences at https://ir.psych.ac.cn/handle/311026/50637.

